# Sensitive, non-invasive detection of chronic wasting disease in wild and captive white-tailed deer using fecal volatile profiling

**DOI:** 10.1101/2025.01.16.633338

**Authors:** Amalia Z. Berna, Tzvi Y. Pollock, Yang Liu, Michelle Gibison, Amritha Mallikarjun, Joey Logan, Cynthia M. Otto, Audrey R. Odom John

## Abstract

Chronic Wasting Disease (CWD) is a universally fatal, transmissible prion disease affecting cervids. Primarily found among deer populations in North America, the disease has spread across the continent and made forays into Europe and Asia as well. Currently, accurate methods for detecting CWD infection require post-mortem dissection of the lymph nodes and brainstem of affected animals. New, high-sensitivity methods of detecting CWD in living animals are sorely needed to help curb spread of this devastating disease in farmed and wild deer. Here, we use two-dimensional gas chromatography-mass spectrometry (GCxGC-MS) to detect volatile organic compounds (VOCs) released from the feces of white-tailed deer (WTD) for differentiation of the feces of CWD-negative and CWD-positive animals. We report four discrete VOCs in captive WTD and ten discrete VOCs in wild WTD, with which we can discriminate CWD-positive and CWD-negative samples. Additionally, we evaluate the ability to detect CWD early during disease progression, by comparing samples from the early stage of infection with samples from late stage and uninfected WTD. Our data suggest that detection of VOCs from the feces of WTD — both in captive and wild populations — can serve as a highly sensitive and non-invasive technique for identifying CWD infection in living animals.

## Introduction

Chronic Wasting Disease (CWD) is a transmissible spongiform encephalopathy of cervids. The etiological agent of CWD is the misfolded form of the prion protein (PrP^CWD^), which associates with the properly folded form of the protein in the brain and triggers a conformational change [1, 2]. CWD is uniformly fatal and can spread rapidly to devastate herds [3]. Infectious prions have been found in the peripheral tissues of infected animals, as well as detected in excreta such as feces, urine, and blood [4–7].

There is no treatment for CWD, and removal from the environment is difficult, as the misfolded protein is resistant to decontamination [8–10]. Additionally, a low inoculum of PrP^CWD^ appears sufficient for oral transmission, leading to rapid spread [5]. As such, prompt detection is critical to disease control. Unfortunately, the standard method for diagnosis of CWD relies on necropsy and evaluation of the retropharyngeal lymph nodes (RPLNs) and brain stem for PrP^CWD^ aggregates by immunohistochemistry (IHC) or enzyme-linked immunosorbent assay (ELISA) [11].

Newer research methods like real-time quaking-induced conversion (RT-QuIC) require samples to be collected via tissue biopsy for best results [4, 12]. The lack of non-invasive diagnostic techniques in living animals has proven an obstacle to disease monitoring and control, especially early during disease progression. As CWD continues to spread, the dire need for antemortem and environmental screening is clear.

Recently, volatile organic compounds (VOCs) have been attractive candidates for minimally invasive disease biomarkers. VOCs are produced by all living creatures as communicative signals or metabolic products. The detection of VOCs in human breath, using methods such gas chromatography/mass spectrometry (GC-MS) or electronic nose systems, has revealed distinct volatiles associated with human diseases such as tuberculosis, malaria, and liver disease [13–17]. In livestock, VOC detection in breath and feces has allowed diagnosis of diseases such as bovine tuberculosis [18]. Studies suggest that fecal samples from captive white-tailed deer (WTD) possess unique VOC profiles based on CWD-positivity [19], and that trained detection dogs can able to differentiate between feces from CWD-positive and CWD-negative deer [20].

In this study, we aimed to use two-dimensional GCxGC-MS VOC detection methods to further characterize the volatile profile of CWD-associated fecal odors, in both wild and captive deer populations. Given the difficulty of diagnosing CWD early during infection, we also sought to evaluate whether VOCs could discriminate early-stage disease. We find that a small handful of VOCs distinguish CWD-positive and -negative samples with good accuracy, both in wild and captive animals. Additionally, our results support the potential for fecal volatiles to identify early-stage CWD-infected animals in the wild.

## Results

### Four volatile compounds discriminate CWD-positive cervids in captivity

A previous study conducted using 1-dimensional GC-MS distinguished feces from CWD-positive and -negative WTD in captive herds using a set of seven VOCs [19]. We sought to utilize our 2-dimensional GCxGC-MS system to further define any CWD-distinguishing fecal volatiles. Fecal samples were collected postmortem from captive animals confirmed by IHC of the RPLNs and brainstem to be either CWD-positive or -negative (Figure 1). VOCs arising from 31 fecal samples (N= 14 positive, N=17 negative) were analyzed through solid phase microextraction (SPME) and GCxGC-MS (Figure 2). Subsequent use of a feature selection algorithm identified four volatile organic compounds as discriminant markers of CWD-positivity, shown in Table 1.

**Figure 1.**
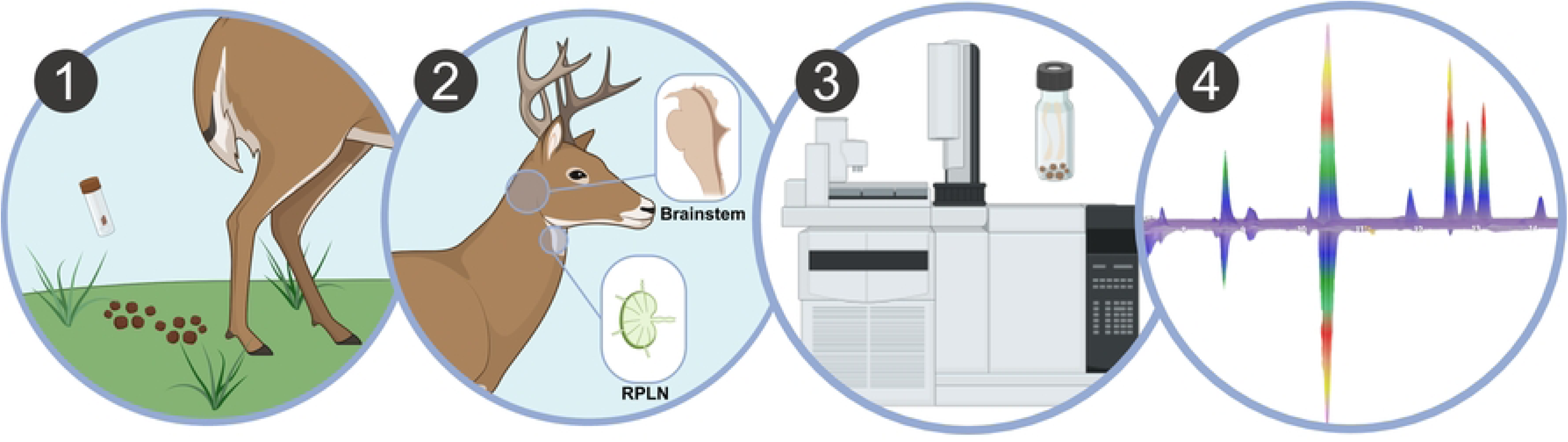
**Fecal samples from white-tail deer are tested for CWD-associated volatiles by GCxGC-MS.** 1.) Fecal samples are removed postmortem from captive or wild WTD. 2.) Animals were classified as CWD-positive or -negative by diagnostic IHC of brainstem and retropharyngeal lymph nodes (RPLN). 3.) Headspace solid-phase microextraction and GCxGC-MS analysis were performed on fecal samples. 4.) Untargeted data analysis identified volatile compounds with discriminatory potential between CWD-positive and -negative samples.

**Figure 2.**
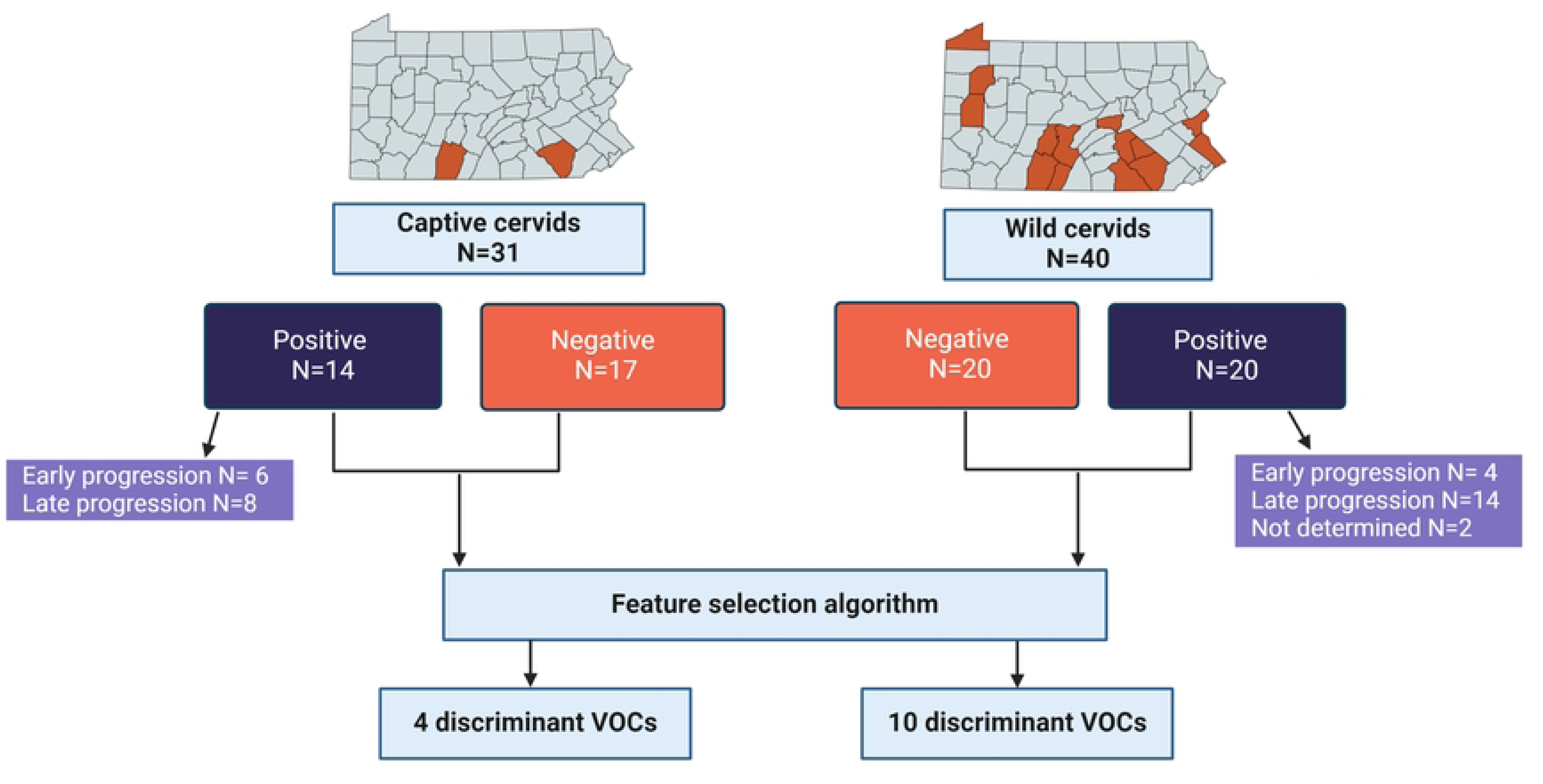
**Sources and infection status of animals tested.** Captive and wild WTD were sampled from the highlighted counties of Pennsylvania, USA. Animals tested were determined to be CWD-positive or -negative as indicated, and CWD-positive animals were characterized as early-stage when PrP^CWD^ was detected in the RPLN and late stage CWD when PrP^CWD^ was detected in both RPLN and brainstem. Positive and negative samples were both used in feature selection, which found discriminant VOCs as shown.

**Table 1:**
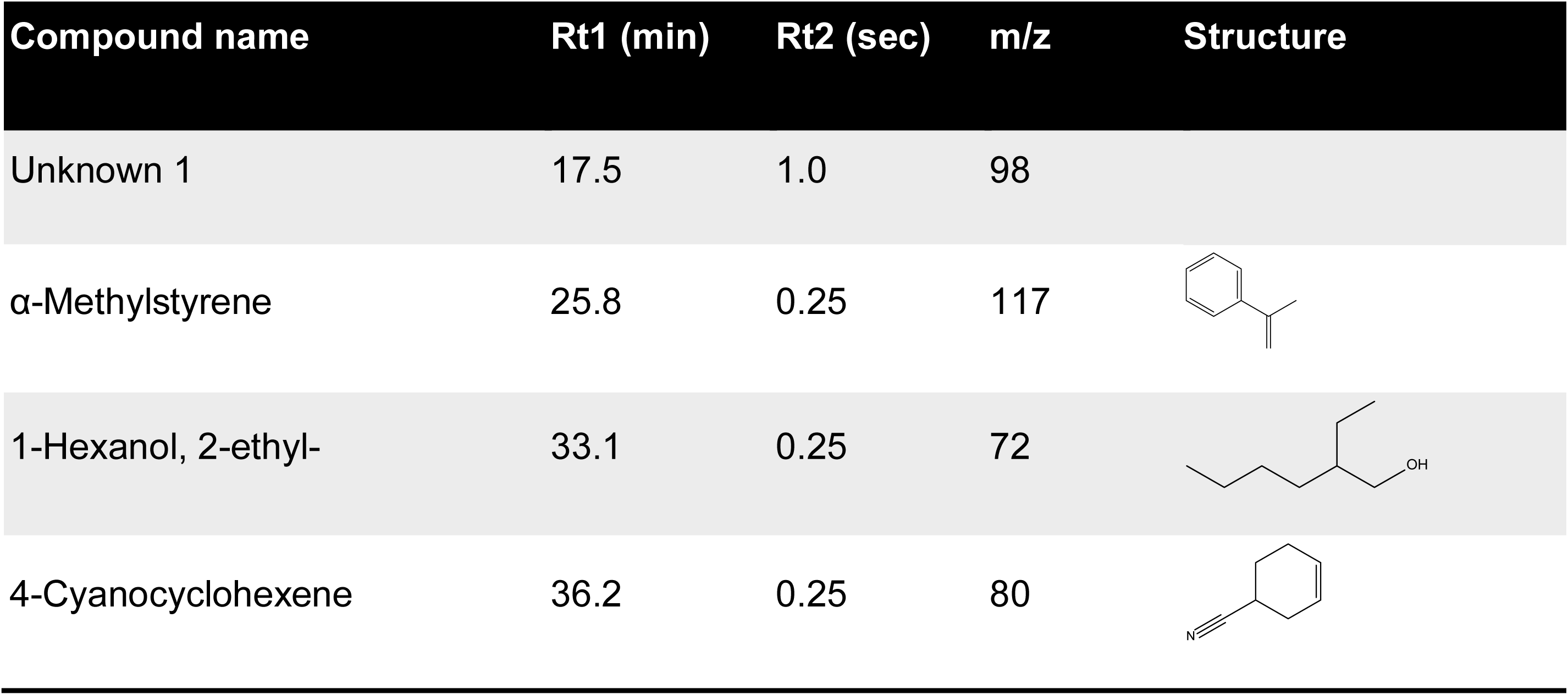
Volatile identification of discriminant features found in fecal matter from captive cervids positive and negative for CWD. Rt1 and Rt2 indicate the retention time for first dimension column D1 (min) and second dimension column D2 (sec) respectively and extracted m/z used for data analysis.

| Compound name | Rt1 (min) | Rt2 (sec) | m/z | Structure |
| --- | --- | --- | --- | --- |
| Unknown 1 | 17.5 | 1.0 | 98 |  |
| α-Methylstyrene | 25.8 | 0.25 | 117 | |
| 1-Hexanol, 2-ethyl- | 33.1 | 0.25 | 72 |  |
| 4-Cyanocyclohexene | 36.2 | 0.25 | 80 |  |

Directly comparing levels of these volatiles revealed a significantly greater abundance of all four compounds in CWD-positive cervids (Figure 3A). Principal component analysis (PCA) revealed distinct clusters of CWD-positive and CWD-negative samples (Figure 3B). Random Forest modeling with Gini Importance measurement identified the four VOCs in Table 1 as potentially discriminant volatiles (Figure S1A), yielding a sensitivity of 57% and a specificity of 82%, for an overall accuracy of 71% (Figure 3C). One of these discriminant volatiles, 2-ethyl-1-hexanol, distinguished CWD infection on its own with a sensitivity of 94% and a specificity of 64% (Figure S1B). Overall, our model boasted moderate accuracy but high specificity, suggesting fecal VOCs might be best suited for screening purposes [21].

**Figure 3.**
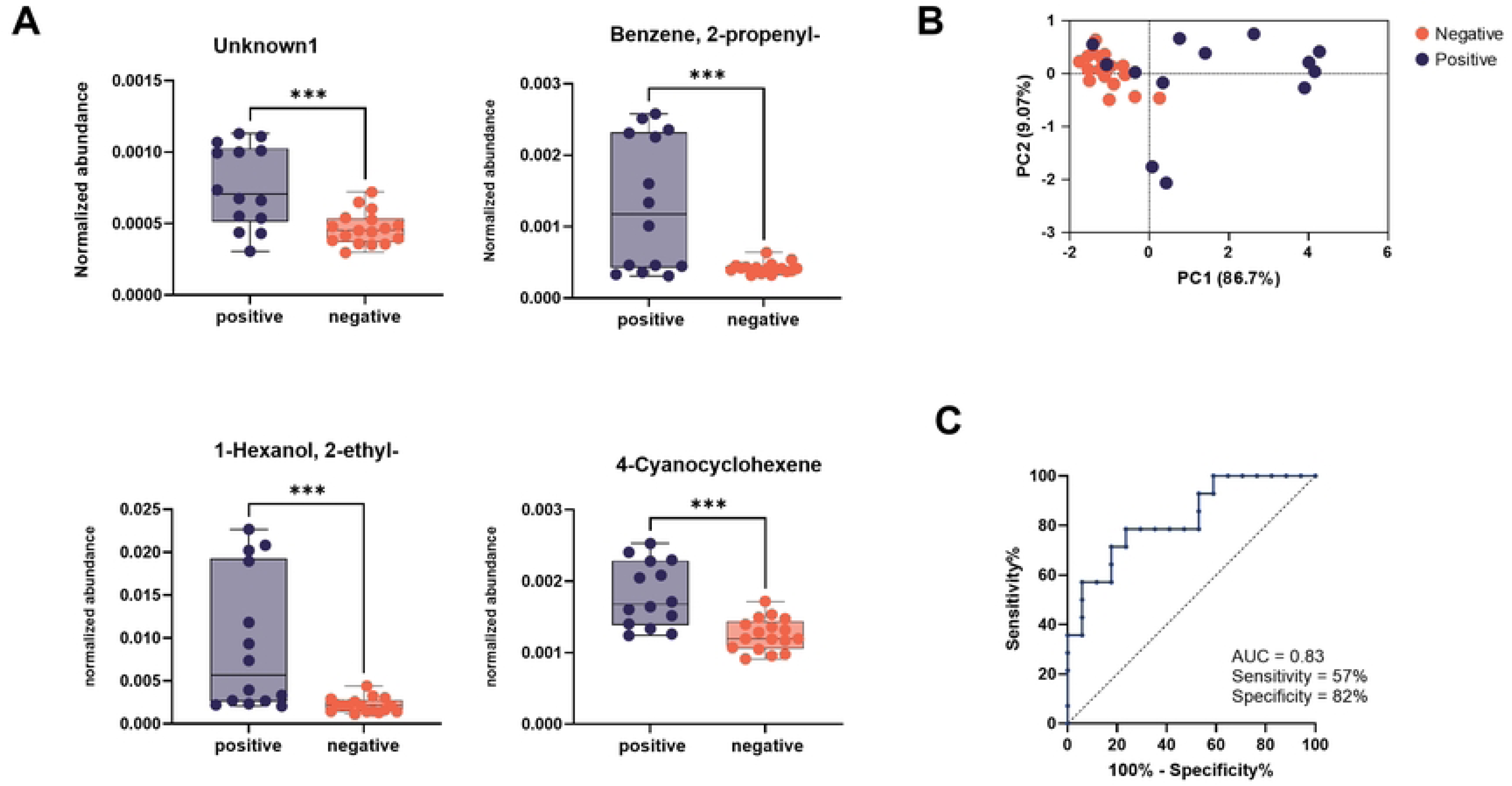
**Four VOCs discriminate between CWD-positive and -negative fecal samples from captive WTD.** A.) Abundances of four volatiles identified during feature selection were compared between positive (N = 14) and negative (N = 17) samples. Abundances were normalized to internal standard compounds. B.) Principal components analysis visualizing the distances between positive and negative fecal sample volatile profiles. C.) Receiver operating characteristic (ROC) curve plotting true vs false positive rate using the four discriminant volatiles to predict infection status. **** p≤ 0.001

Asymptomatic animals can spread infectious PrP^CWD^ in the environment, and so minimally invasive diagnostic techniques that can detect early-stage infection are required. To this end, we compared early-and late-stage CWD. Presence of PrP^CWD^ in the RPLNs alone was defined as early-stage, while presence in both RPLNs and brainstem was defined as late-stage [22]. Our preliminary analysis revealed significantly decreased levels of the VOCs 2-ethyl-1-hexanol and 2-propenyl-benzene in early-stage, compared to late-stage CWD (Figure S2A-B). However, no clustering was observed between early-stage CWD-infected captive cervids and healthy controls by PCA (Figure S2C). Since our sample size was small, this preliminary analysis is underpowered, and future studies will require a larger population of early-stage CWD.

Finally, we manually searched for a subset of the volatiles previously found to be discriminant [19]. No significant differences in these compounds were detected between CWD-positive and CWD-negative samples (Figure S3A).

### Ten volatile compounds discriminate CWD-positive cervids in field settings

VOC-based detection of CWD infection has previously focused on farmed WTD, however CWD infects both farmed and free-ranging populations [3]. We therefore used our proven GCxGC-MS system to evaluate whether fecal VOCs might also discriminate between CWD-positive and CWD-negative animals in field settings. Forty fecal samples (N=20 positive, N=20 negative) were procured from wild WTD collected from hunter harvests, roadkill, or clinical suspects (Figure 2). Animals were confirmed to be CWD-positive or CWD-negative by repeated ELISA and confirmatory IHC (Figure 1) [11]. Fecal samples were subjected to SPME isolation of VOCs, followed by GCxGC-MS analysis. Feature discovery identified ten VOCs as potentially discriminant features of CWD-positivity, shown in Table 2.

**Table 2:**
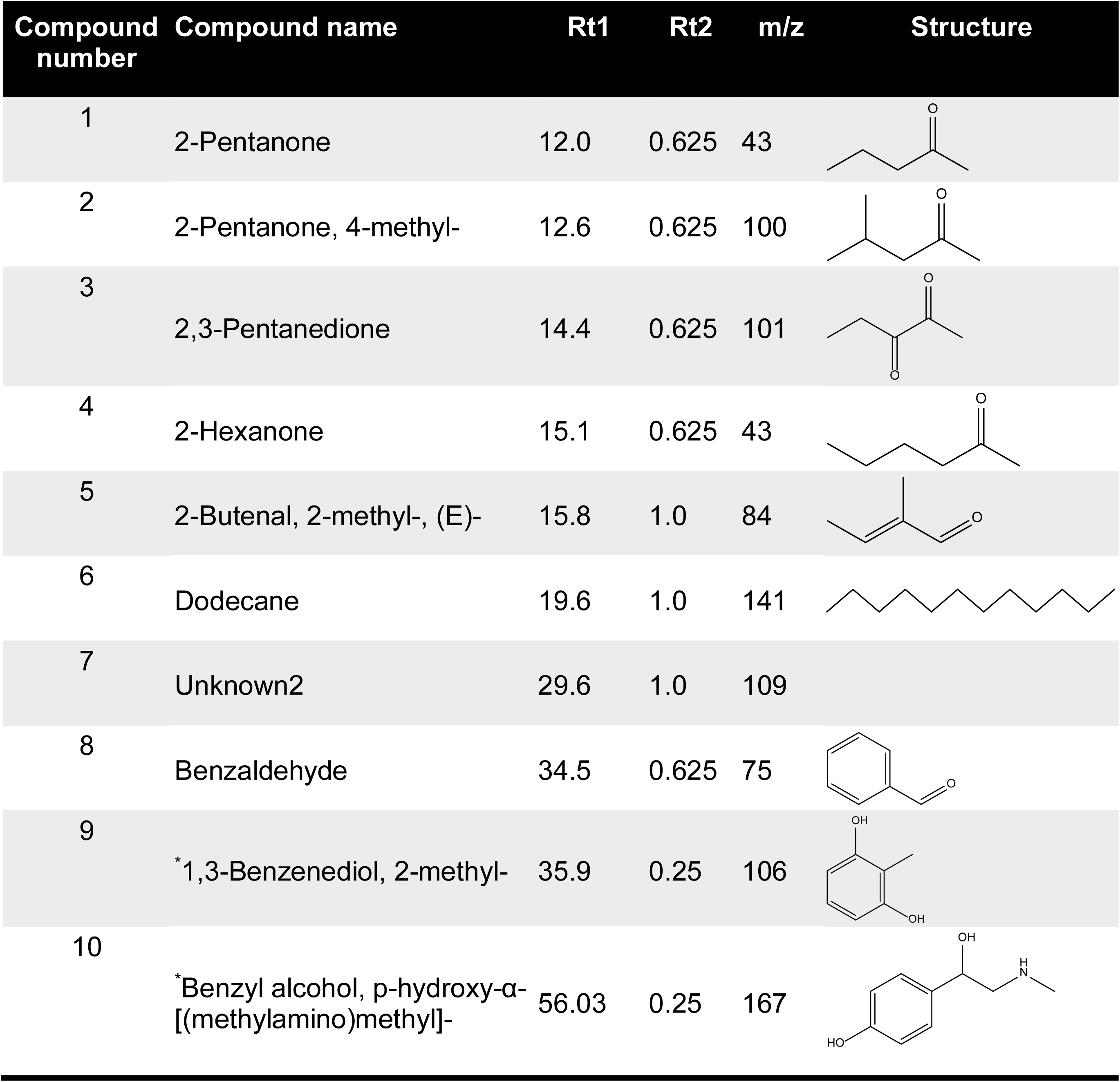
Volatile identification of discriminant features found in fecal matter from wild-type cervids positive and negative for CWD. Rt1 and Rt2 indicated the retention time for first dimension column D1 (min) and second dimension column D2 (sec) respectively and extracted m/z used for data analysis.

| Compound number | Compound name | Rt1 | Rt2 | m/z | Structure |
| --- | --- | --- | --- | --- | --- |
| 1 | 2-Pentanone | 12.0 | 0.625 | 43 |  |
| 2 | 2-Pentanone, 4-methyl- | 12.6 | 0.625 | 100 |  |
| 3 | 2,3-Pentanedione | 14.4 | 0.625 | 101 |  |
| 4 | 2-Hexanone | 15.1 | 0.625 | 43 |  |
| 5 | 2-Butenal, 2-methyl-, (E)- | 15.8 | 1.0 | 84 |  |
| 6 | Dodecane | 19.6 | 1.0 | 141 |  |
| 7 | Unknown2 | 29.6 | 1.0 | 109 |  |
| 8 | Benzaldehyde | 34.5 | 0.625 | 75 |  |
| 9 | *1,3-Benzenediol, 2-methyl- | 35.9 | 0.25 | 106 |  |
| 10 | *Benzyl alcohol, p-hydroxy-α-[(methylamino)methyl]- | 56.03 | 0.25 | 167 |  |

Four of the compounds isolated in our screen were lower in the feces of CWD-positive animals compared to CWD-negative animals, while the reverse was true of the other six compounds (Figure 4A). PCA revealed that wild CWD-positive and CWD-negative samples formed two distinct clusters, indicating substantial differences in their VOC profiles (Figure 4B). Random Forest modeling using these ten discriminant VOCs resulted in 85% sensitivity and 90% specificity, for an accuracy of 88% (Figure 4C). These data suggest that fecal volatile profiling has strong potential as a non-invasive, highly accurate tool for detection of CWD in wild populations.

**Figure 4.**
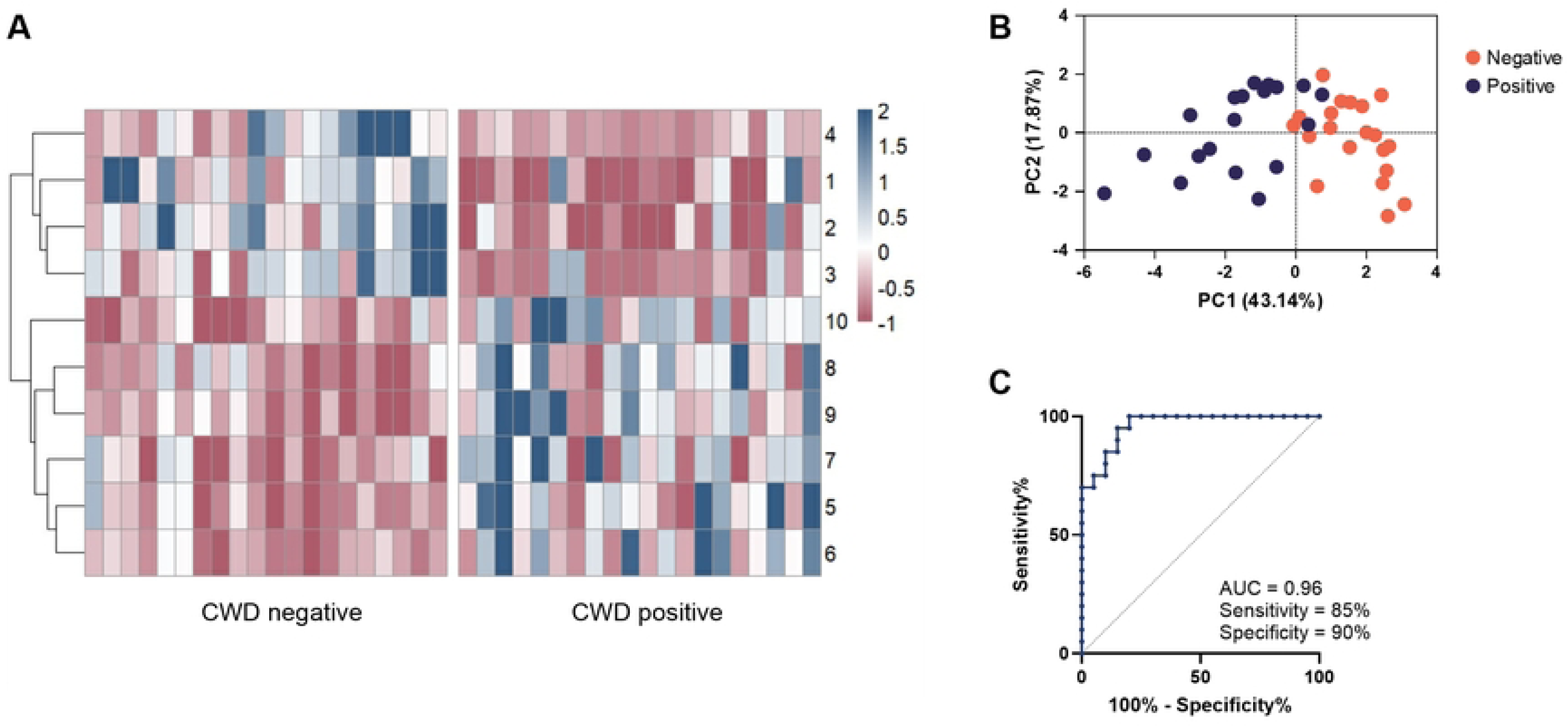
**Ten VOCs discriminate between CWD-positive and -negative fecal samples from wild WTD.** A.) *z* score abundances of ten discriminant volatiles in CWD-negative (N = 20) and CWD-positive (N = 20) fecal samples. Compound number correlates to the respective compound in Table B.) Principal components analysis visualizing the distances between positive and negative fecal sample volatile profiles. C.) Receiver operating characteristic (ROC) curve plotting true vs false positive rate using the ten discriminant volatiles to predict infection status.

Much like in captive cervids, early detection of CWD in wild cervid populations is crucial to combatting spread. Our preliminary analysis demonstrated no separation between the VOC profiles of early and late-stage CWD among wild cervids (Figure S5A). As we observed that levels of 2-ethyl-1-hexanol and 2-propenyl benzene were elevated in late-stage CWD among captive animals (Figure S2A-B), we compared levels of these two volatiles in wild samples as well. We observed no separation between early and late-stage CWD in wild cervids (Figure S5B). Further-more, PCA revealed that early-stage CWD samples cluster away from healthy controls (Figure S5C). These data suggest that fecal VOC collection can detect even early-stage CWD in wild cervids. As with our data in captive cervids, however, the small sample size of early-stage CWD (N = 4) is a major limitation, and further studies are required.

We again searched for a subset of the discriminant volatiles identified in a previous study of captive WTD [19], finding that 1-butanol levels were increased in feces from CWD-negative deer, while 3-methyl-indole levels were increased in feces from CWD-positive deer (Figure S3B).

## Discussion

The deer prion disease CWD has no known treatments and is uniformly fatal. A small inoculum of PrP^CWD^ is sufficient to transmit orally, leading to rapid spread through herds of WTD [5]. Furthermore, animals can shed infectious prions long before symptoms [6]. The need for new forms of minimally invasive, antemortem testing for CWD is clear and pressing [3]. Diagnosis of infection using VOC profiling is a growing field, with studies detecting infections such as tuberculosis and malaria, among others [17, 23]. We demonstrate in this study that fecal volatiles accurately differentiate feces from CWD-positive and CWD-negative deer. Our work demonstrates the promise for fecal VOC analysis as a method of CWD surveillance in captive and wild animal populations.

Our study reveals four volatile compounds, identified in Table 1, which discriminated CWD infection in captive WTD with a classification accuracy of 71% (Figure 3). Likewise, our study reveals 10 volatile compounds, identified in Table 2, which discriminated CWD infection in wild WTD with a classification accuracy of 88% (Figure 4). None of the identified VOCs are known to have eukaryotic origin, and are postulated to originate from diet or the microbiome. The VOCs identified in this study did not overlap with those seen previously [19]; differences between the volatiles found here and in previous work may derive from our more sensitive instrumentation, as well as from natural variation between populations of captive and wild cervids. Further validation of these VOC profiles in diverse cervid populations is needed. However, our work here represents a strong framework for future diagnostic testing.

Perhaps surprisingly, the fecal VOCs that proved discriminant for captive WTD and wild WTD did not feature any common compounds. Additionally, fecal VOC profiles of wild WTD demonstrated far greater sensitivity than captive WTD, with 85% sensitivity for the former and 57% sensitivity for the latter (Figure 3C, 4C). One possible cause for these discrepancies lies in the effect of captivity on the intestinal microbiome, a key biological source of fecal VOCs. Previous studies have shown that captivity alone impacts the fecal microbiota of cervids [24, 25], and that CWD infection can further alter the microbiome [24, 26]. Another possible explanation lies in the dietary habits of infected cervids. Research shows that infected cervids struggle to maintain efficient foraging activity [6, 27]. This difficulty foraging may prove far less impactful in captive WTD, provided ample food by farmers, than in wild populations, explaining the greater discrimination observed by fecal volatile analysis.

Detecting early-stage CWD, ideally prior to symptom onset, would be optimal for long-term control of spread. While further targeted investigation is required, our early data demonstrate that VOCs during early-stage CWD bear similarity to late-stage CWD. This is especially apparent in samples from wild cervids (Figure S5). These data suggest that fecal VOC analysis is a promising method for early detection of CWD in both captive and wild cervids. Following further optimization, non-invasive detection of CWD-associated volatiles *in situ* among WTD populations could greatly empower public health surveillance for CWD. Modern technologies such as electronic nose (“e-nose”) sensor arrays [23] or miniature mass spectrometers could further enhance the speed and efficiency with which CWD is detected and its spread limited.

---

**Figure S1.**
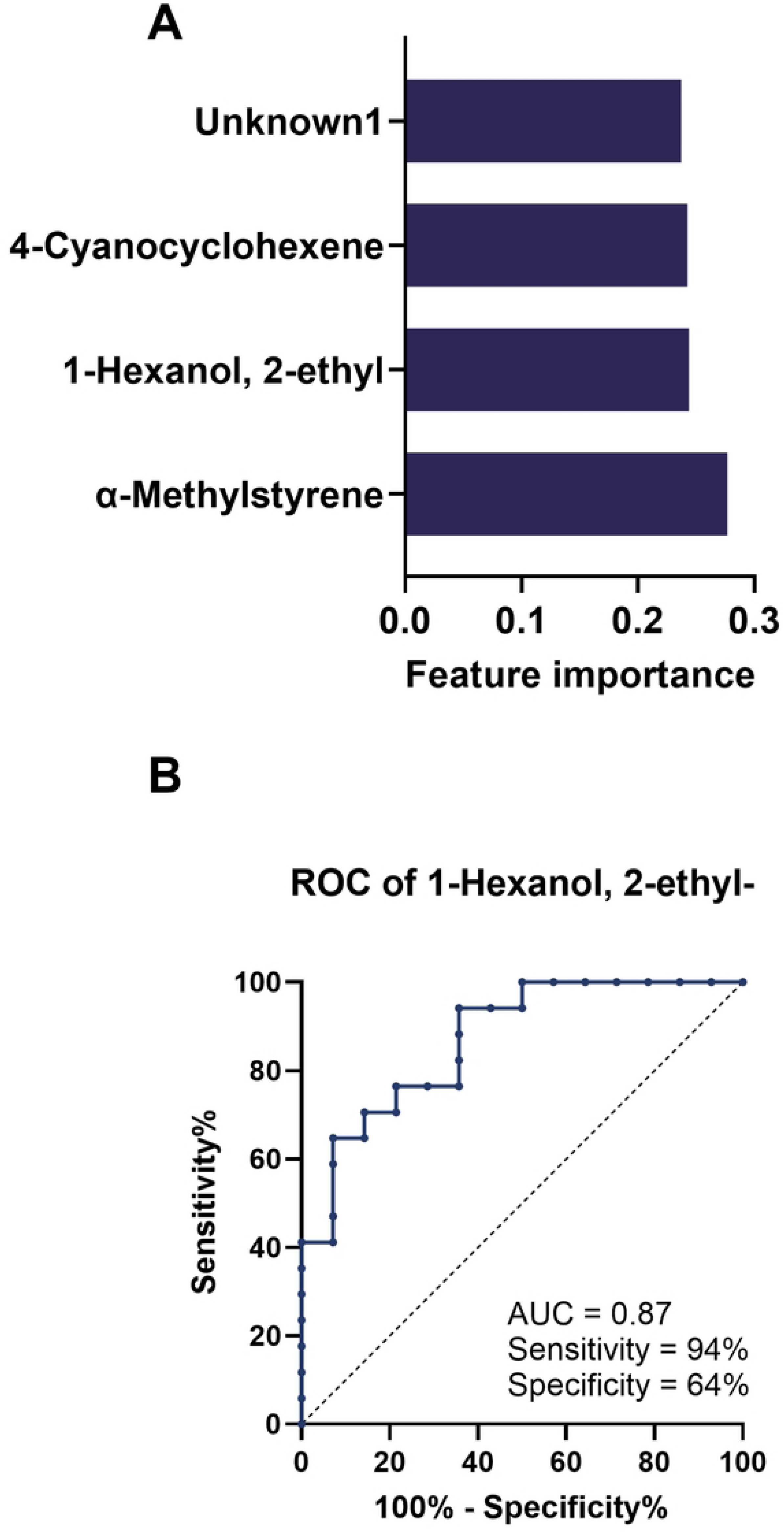
**Volatiles discovered in captive WTD.** A.) Feature importance of the four discriminant volatiles was determined using Random Forest Classifier with Gini importance. B.) Receiver operating characteristic (ROC) curve plotting true vs false positive rate using 1-Hexanol, 2-ethyl alone to predict infection status.

**Figure S2.**
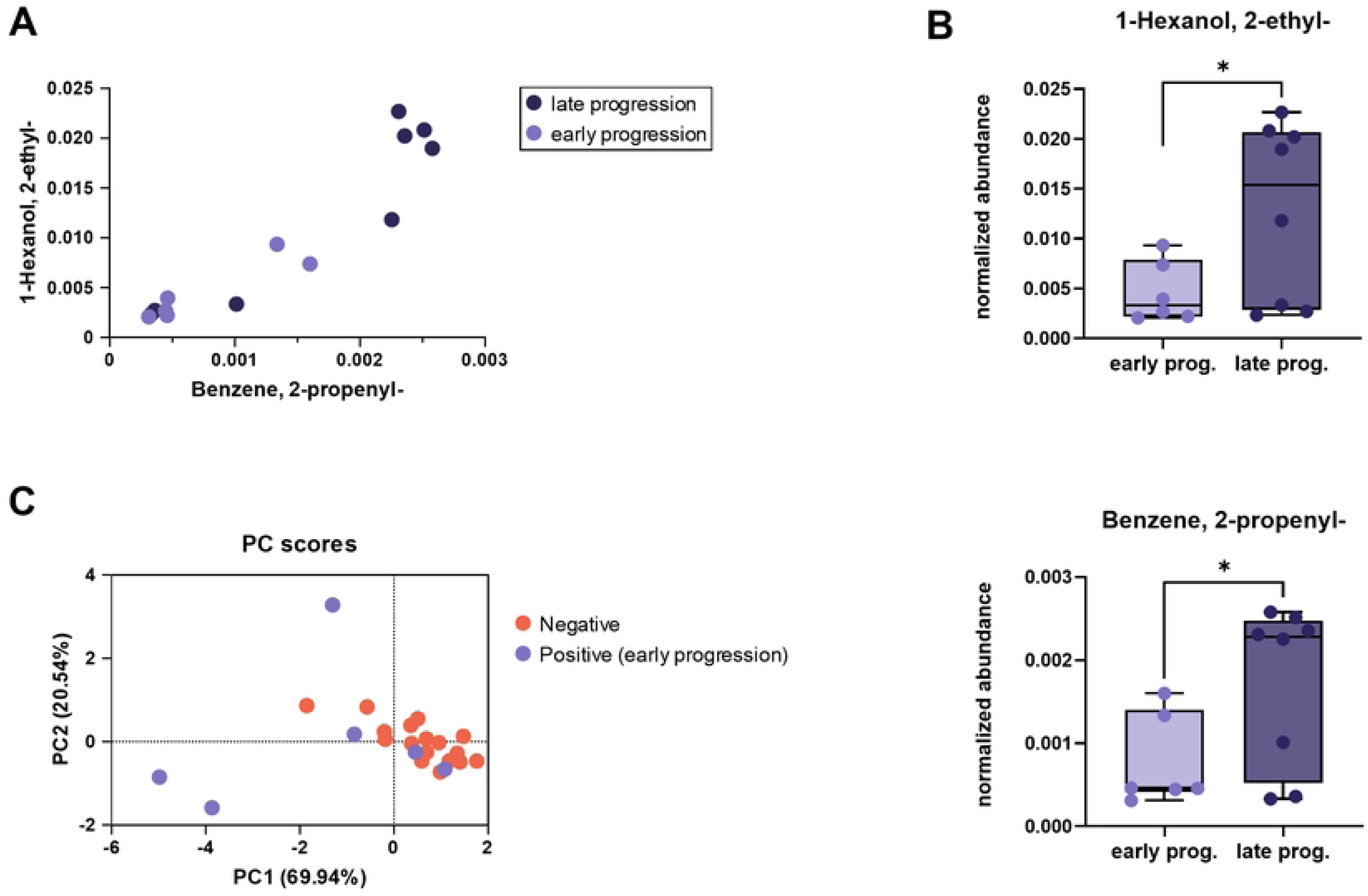
**Early and late progression CWD differ in VOC abundance in captive WTD.** A-B.) The relative abundances of both 1-Hexanol, 2-ethyl and Benzene, 2-propenyl during early (N = 6) and late (N = 8) stage CWD infection are compared. Abundances are normalized to internal standard control. C.) Principal components analysis visualizing the distances between early stage positive and negative fecal sample volatile profiles. * p ≤ 0.05

**Figure S3.**
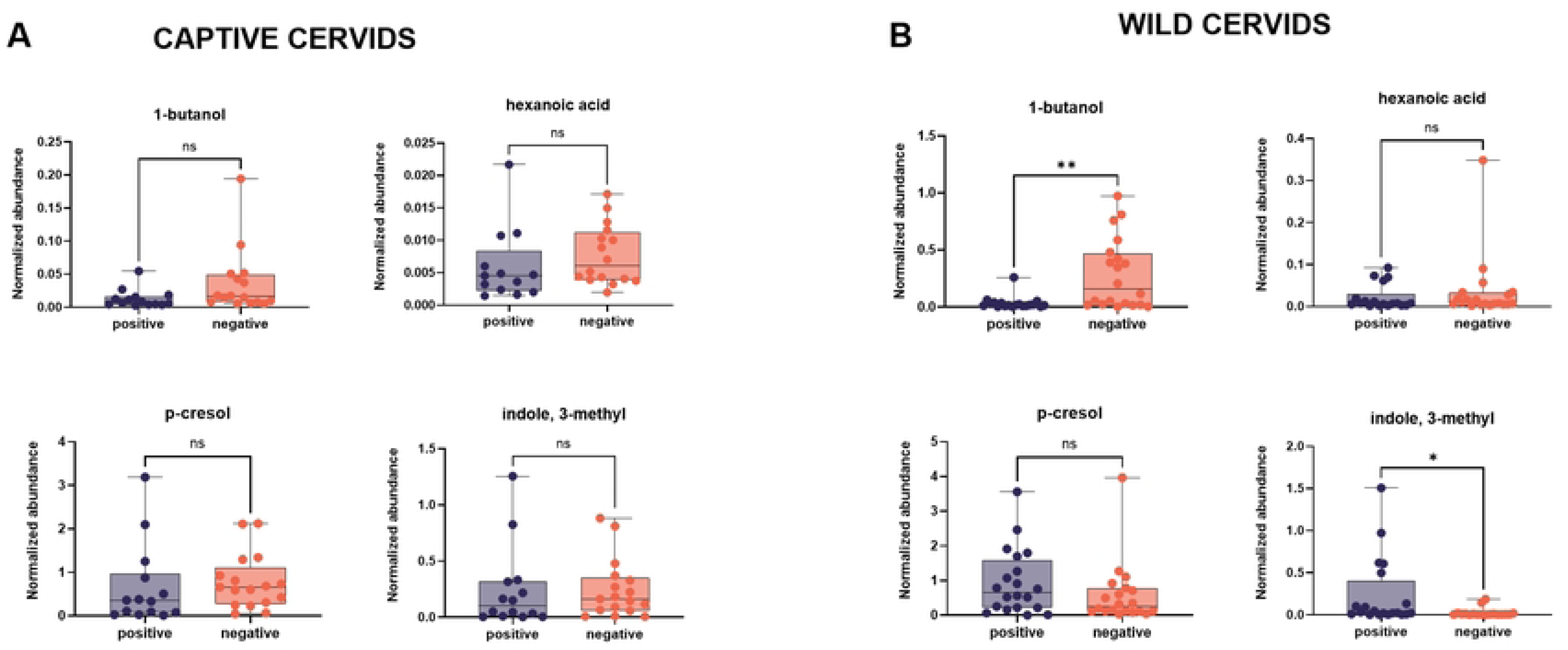
Previously described volatiles do not discriminate between CWD positivity in fecal samples. Abundances of 1-butanol; hexanoic acid; p-cresol; and indole, 3-methyl were compared between positive and negative fecal samples from captive (A) or wild (B) WTD. Abundances were normalized to internal standard control. ns = no significance, * p ≤ 0.05.

**Figure S4.**
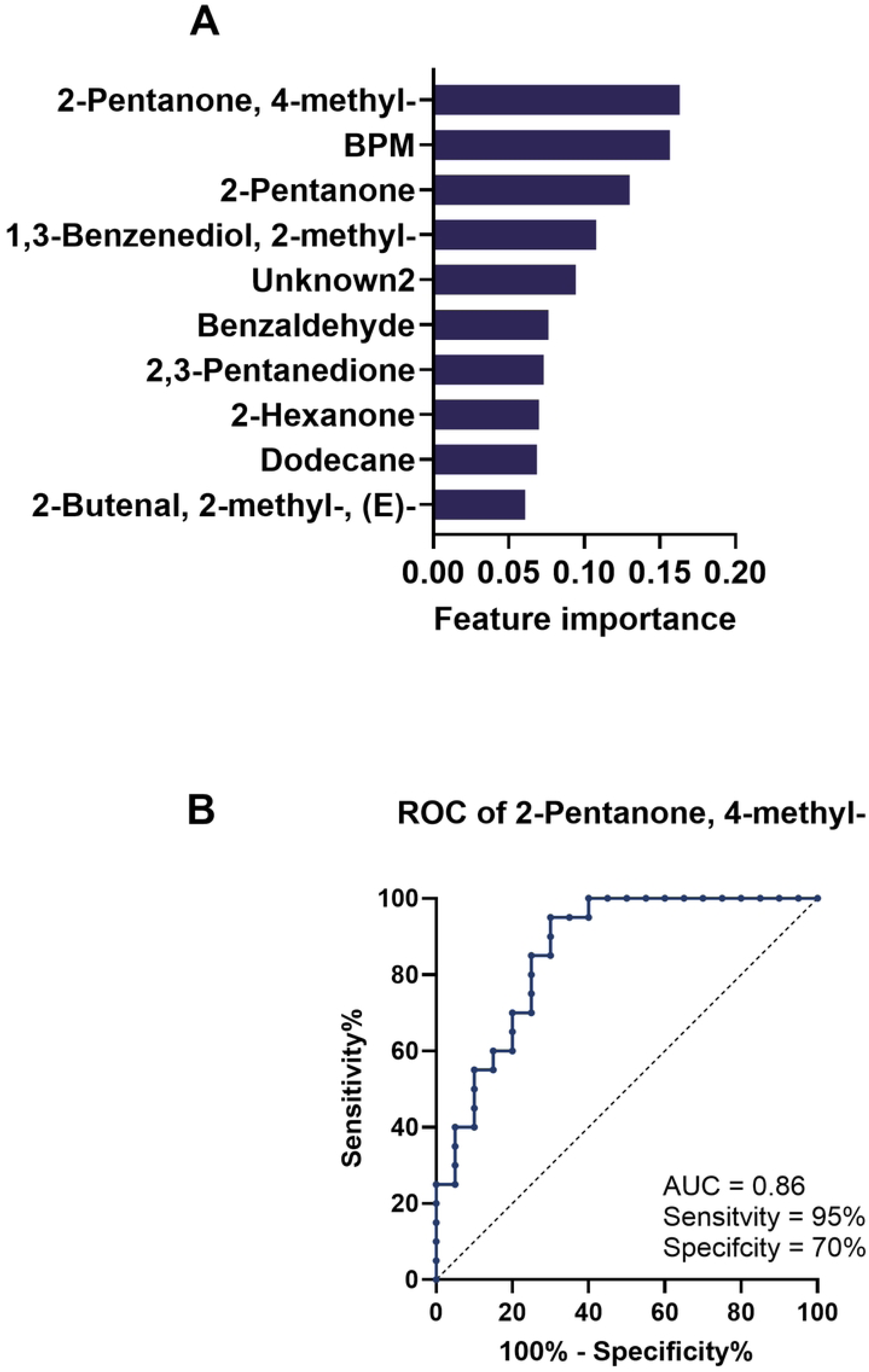

**Figure S5.**
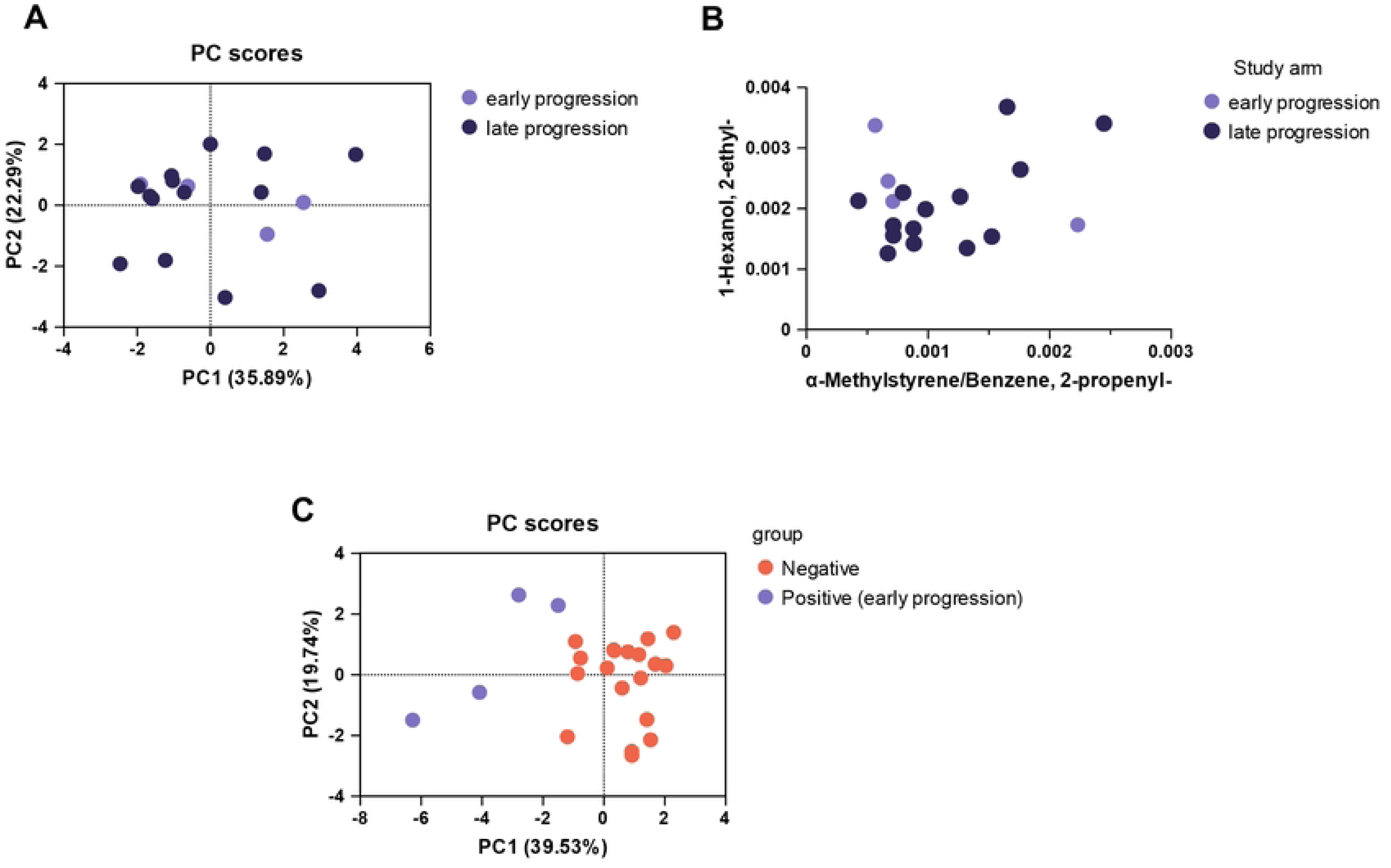
**Fecal samples from early progression CWD can be discriminated from CWD-negative samples in wild WTD.** A.) Principal components analysis visualizing the distances between early (N = 4) and late (N = 14) stage CWD fecal sample volatile profiles B.) The relative abundances of both 1-Hexanol, 2-ethyl and Benzene, 2-propenyl during early (N = 4) and late (N = 14) progression CWD infection are compared. Abundances are normalized to internal standard control. C.) Principal components analysis visualizing the distances between early progression positive and negative fecal sample volatile profiles.

## References

1. Williams ES, Young S. Chronic wasting disease of captive mule deer: a spongiform encephalopathy. J Wildl Dis. 1980;16(1):89–98. doi: 10.7589/0090-3558-16.1.89. PubMed PMID: 7373730.

2. Haley NJ, Hoover EA. Chronic wasting disease of cervids: current knowledge and future perspectives. Annu Rev Anim Biosci. 2015;3:305–25. Epub 20141002. doi: 10.1146/annurev-animal-022114-111001. PubMed PMID: 25387112.

3. Mori J, Rivera N, Novakofski J, Mateus-Pinilla N. A review of chronic wasting disease (CWD) spread, surveillance, and control in the United States captive cervid industry. Prion. 2024;18(1):54–67. Epub 20240422. doi: 10.1080/19336896.2024.2343220. PubMed PMID: 38648377; PubMed Central PMCID: PMCPMC11037284.

4. John TR, Schatzl HM, Gilch S. Early detection of chronic wasting disease prions in urine of pre-symptomatic deer by real-time quaking-induced conversion assay. Prion. 2013;7(3):253–8. doi: 10.4161/pri.24430. PubMed PMID: 23764839; PubMed Central PMCID: PMCPMC3783112.

5. Denkers ND, Hoover CE, Davenport KA, Henderson DM, McNulty EE, Nalls AV, et al. Very low oral exposure to prions of brain or saliva origin can transmit chronic wasting disease. PLoS One. 2020;15(8):e0237410. Epub 20200820. doi: 10.1371/journal.pone.0237410. PubMed PMID: 32817706; PubMed Central PMCID: PMCPMC7446902.

6. Henderson DM, Denkers ND, Hoover CE, Garbino N, Mathiason CK, Hoover EA. Longitudinal Detection of Prion Shedding in Saliva and Urine by Chronic Wasting Disease-Infected Deer by Real-Time Quaking-Induced Conversion. J Virol. 2015;89(18):9338–47. Epub 20150701. doi: 10.1128/JVI.01118-15. PubMed PMID: 26136567; PubMed Central PMCID: PMCPMC4542351.

7. Pulford B, Spraker TR, Wyckoff AC, Meyerett C, Bender H, Ferguson A, et al. Detection of PrPCWD in feces from naturally exposed Rocky Mountain elk (*Cervus elaphus nelsoni*) using protein misfolding cyclic amplification. J Wildl Dis. 2012;48(2):425–34. doi: 10.7589/0090-3558-48.2.425. PubMed PMID: 22493117.

8. Taylor D. Inactivation of the causal agents of Creutzfeldt-Jakob disease and other human prion diseases. Brain Pathol. 1996;6(2):197–8. doi: 10.1111/j.1750-3639.1996.tb00800.x. PubMed PMID: 8737933.

9. Sehulster LM. Prion inactivation and medical instrument reprocessing: challenges facing healthcare facilities. Infect Control Hosp Epidemiol. 2004;25(4):276–9. doi: 10.1086/502391. PubMed PMID: 15108722.

10. Williams K, Hughson AG, Chesebro B, Race B. Inactivation of chronic wasting disease prions using sodium hypochlorite. PLoS One. 2019;14(10):e0223659. Epub 20191004. doi: 10.1371/journal.pone.0223659. PubMed PMID: 31584997; PubMed Central PMCID: PMCPMC6777796.

11. Hibler CP, Wilson KL, Spraker TR, Miller MW, Zink RR, DeBuse LL, et al. Field validation and assessment of an enzyme-linked immunosorbent assay for detecting chronic wasting disease in mule deer (*Odocoileus hemionus*), white-tailed deer (*Odocoileus virginianus*), and Rocky Mountain elk (*Cervus elaphus nelsoni*). J Vet Diagn Invest. 2003;15(4):311–9. doi: 10.1177/104063870301500402. PubMed PMID: 12918810.

12. Piel RB, 3rd, Veneziano SE, Nicholson EM, Walsh DP, Lomax AD, Nichols TA, et al. Validation of a real-time quaking-induced conversion (RT-QuIC) assay protocol to detect chronic wasting disease using rectal mucosa of naturally infected, pre-clinical white-tailed deer (*Odocoileus virginianus*). PLoS One. 2024;19(6):e0303037. Epub 20240613. doi: 10.1371/journal.pone.0303037. PubMed PMID: 38870153; PubMed Central PMCID: PMCPMC11175469.

13. Pollock TY, Odom John AR. Thinking Small, Stinking Big: The World of Microbial Odors. J Infect Dis. 2024;229(5):1254–5. doi: 10.1093/infdis/jiad405. PubMed PMID: 37738417; PubMed Central PMCID: PMCPMC11095540.

14. Berna AZ, Akaho EH, Harris RM, Congdon M, Korn E, Neher S, et al. Reproducible breath metabolite changes in children with SARS-CoV-2 infection. medRxiv. 2021. Epub 20210507. doi: 10.1101/2020.12.04.20230755. PubMed PMID: 33330891; PubMed Central PMCID: PMCPMC7743102.

15. Berna AZ, Odom John AR. Breath Metabolites to Diagnose Infection. Clin Chem. 2021;68(1):43–51. doi: 10.1093/clinchem/hvab218. PubMed PMID: 34969107; PubMed Central PMCID: PMCPMC8718131.

16. Kelly M, Su CY, Schaber C, Crowley JR, Hsu FF, Carlson JR, et al. Malaria parasites produce volatile mosquito attractants. mBio. 2015;6(2). Epub 20150324. doi: 10.1128/mBio.00235-15. PubMed PMID: 25805727; PubMed Central PMCID: PMCPMC4453533.

17. Schaber CL, Katta N, Bollinger LB, Mwale M, Mlotha-Mitole R, Trehan I, et al. Breathprinting Reveals Malaria-Associated Biomarkers and Mosquito Attractants. J Infect Dis. 2018;217(10):1553–60. doi: 10.1093/infdis/jiy072. PubMed PMID: 29415208; PubMed Central PMCID: PMCPMC6279169.

18. Brebu M, Simion VE, Andronie V, Jaimes-Mogollon AL, Beleno-Saenz KJ, Ionescu F, et al. Putative volatile biomarkers of bovine tuberculosis infection in breath, skin and feces of cattle. Mol Cell Biochem. 2023;478(11):2473–80. Epub 20230225. doi: 10.1007/s11010-023-04676-5. PubMed PMID: 36840799.

19. Ellis CK, Volker SF, Griffin DL, VerCauteren KC, Nichols TA. Use of faecal volatile organic compound analysis for ante-mortem discrimination between CWD-positive, -negative exposed, and -known negative white-tailed deer (*Odocoileus virginianus*). Prion. 2019;13(1):94–105. doi: 10.1080/19336896.2019.1607462. PubMed PMID: 31032718; PubMed Central PMCID: PMCPMC7000150.

20. Mallikarjun A, Swartz B, Kane SA, Gibison M, Wilson I, Collins A, et al. Canine detection of chronic wasting disease (CWD) in laboratory and field settings. Prion. 2023;17(1):16–28. doi: 10.1080/19336896.2023.2169519. PubMed PMID: 36740856; PubMed Central PMCID: PMCPMC9904315.

21. Simundic AM. Measures of Diagnostic Accuracy: Basic Definitions. EJIFCC. 2009;19(4):203–11. Epub 20090120. PubMed PMID: 27683318; PubMed Central PMCID: PMCPMC4975285.

22. Keane DP, Barr DJ, Keller JE, Hall SM, Langenberg JA, Bochsler PN. Comparison of retropharyngeal lymph node and obex region of the brainstem in detection of chronic wasting disease in white-tailed deer (*Odocoileus virginianus*). J Vet Diagn Invest. 2008;20(1):58–60. doi: 10.1177/104063870802000110. PubMed PMID: 18182509.

23. Coronel Teixeira R, Rodriguez M, Jimenez de Romero N, Bruins M, Gomez R, Yntema JB, et al. The potential of a portable, point-of-care electronic nose to diagnose tuberculosis. J Infect. 2017;75(5):441–7. Epub 20170810. doi: 10.1016/j.jinf.2017.08.003. PubMed PMID: 28804027.

24. Minich D, Madden C, Evans MV, Ballash GA, Barr DJ, Poulsen KP, et al. Alterations in gut microbiota linked to provenance, sex, and chronic wasting disease in white-tailed deer (*Odocoileus virginianus*). Sci Rep. 2021;11(1):13218. Epub 20210624. doi: 10.1038/s41598-021-89896-9. PubMed PMID: 34168170; PubMed Central PMCID: PMCPMC8225879.

25. Pacheco-Torres I, Hernandez-Sanchez D, Garcia-De la Pena C, Tarango-Arambula LA, Crosby-Galvan MM, Sanchez-Santillan P. Analysis of the Intestinal and Faecal Bacterial Microbiota of the Cervidae Family Using 16S Next-Generation Sequencing: A Review. Microorganisms. 2023;11(7). Epub 20230724. doi: 10.3390/microorganisms11071860. PubMed PMID: 37513032; PubMed Central PMCID: PMCPMC10386072.

26. Didier A, Bourner M, Kleks G, Zolty A, Kumar B, Nichols T, et al. Prospective fecal microbiomic biomarkers for chronic wasting disease. Microbiol Spectr. 2024;12(3):e0375022. Epub 20240201. doi: 10.1128/spectrum.03750-22. PubMed PMID: 38299851; PubMed Central PMCID: PMCPMC10913453.

27. Fox KA, Jewell JE, Williams ES, Miller MW. Patterns of PrPCWD accumulation during the course of chronic wasting disease infection in orally inoculated mule deer (*Odocoileus hemionus*). J Gen Virol. 2006;87(Pt 11):3451–61. doi: 10.1099/vir.0.81999-0. PubMed PMID: 17030882.

28. Ferreira NC, Charco JM, Plagenz J, Orru CD, Denkers ND, Metrick MA, 2nd, et al. Detection of chronic wasting disease in mule and white-tailed deer by RT-QuIC analysis of outer ear. Sci Rep. 2021;11(1):7702. Epub 20210408. doi: 10.1038/s41598-021-87295-8. PubMed PMID: 33833330; PubMed Central PMCID: PMCPMC8032746.

29. Spadafora ND, Eggermont D, Krestakova V, Chenet T, Van Rossum F, Purcaro G. Comprehensive analysis of floral scent and fatty acids in nectar of *Silene nutans* through modern analytical gas chromatography techniques. J Chromatogr A. 2023;1696:463977. Epub 20230407. doi: 10.1016/j.chroma.2023.463977. PubMed PMID: 37054636.

